# Mucosal Inflammation as a Driver of Gd-IgA1 Production in IgA Nephropathy: Mechanistic Support for Targeted Therapies

**DOI:** 10.64898/2026.10.19.759132

**Authors:** Katerina Zachova, Alica Cutkova, Petr Kosztyu, Yukako Ohyama, Kazuo Takahashi, Josef Zadrazil, Jiri Orsag, Nadezda Petejova, Dana Galuszkova, Jiri Mestecky, Milan Raska

**Affiliations:** Department of Immunology, Faculty of Medicine and Dentistry, Palacky University Olomouc, Olomouc, Czech Republic; Department of Immunology, University Hospital Olomouc, Olomouc, Czech Republic; Department of Biomedical Molecular Sciences, School of Medicine, Fujita Health University, Nagoya, Aichi, Japan; Department of Internal Medicine III Nephrology, Rheumatology and Endocrinology, University Hospital Olomouc, Olomouc, Czech Republic; Department of Internal Medicine III Nephrology, Rheumatology and Endocrinology, Faculty of Medicine and Dentistry, Palacky University Olomouc, Czech Republic; Department of Internal Medicine and Cardiology University Hospital Ostrava and Faculty of Medicine University of Ostrava, Czech Republic; Department of Transfusion Medicine, University Hospital Olomouc, Olomouc, Czech Republic; Departments of Microbiology and Medicine, University of Alabama at Birmingham, Birmingham, Alabama, USA; Laboratory of Cellular and Molecular Immunology of the Czech Academy of Sciences, Prague, Czech Republic

**Keywords:** IgA nephropathy, polymeric λ-enriched Galactose-deficient IgA1 (Gd-IgA1), Mucosal immune activation, B cell dysregulation, Plasma cells, BAFF/APRIL, Gut–kidney axis, Targeted therapy

## Abstract

**Rationale & Objective:** Immunoglobulin A nephropathy (IgAN) is characterized by mesangial deposition of immune complexes containing galactose-deficient IgA1 (Gd-IgA1), predominantly with λ light chains and Gd-IgA1 glycans-recognizing antibodies. Disease exacerbations are frequently linked to mucosal infections, underscoring the role of dysregulated mucosal immunity. Whereas current therapeutic strategies targeting Gd-IgA1-producing cells are largely evaluated based on clinical endpoints, a critical gap remains in understanding how these therapies influence Gd-IgA production and whether targeting mucosal immunity represents an effective therapeutic strategy. Here, we provide a mechanistic insight into Gd-IgA1+ B cell behavior within inflammatory mucosal-like microenvironment.

**Study Design:** Case–control study with *ex vivo* functional analysis.

**Setting & Participants:** Peripheral blood mononuclear cells from patients with biopsy-proven IgAN and healthy controls.

**Exposure(s):** Stimulation with a cytokine combination mimicking mucosal immune activation (IL-6, IL-21, BAFF, APRIL).

**Outcome(s):** Differentiation of B cell subsets involved in IgAN and production of IgA, Gd-IgA1, and IgG.

**Analytical Approach:** Flow cytometry, ELISA, cytometric bead array, and Western blotting.

**Results:** Mucosal cytokine stimulation induced robust differentiation of B cells into CD38^high^ plasmablasts and CD138^high^ plasma cells, with a disproportionate expansion of Gd-IgA1^+^ populations in IgAN. Compared with controls, PBMCs from IgAN patients demonstrated markedly increased production of Gd-IgA1, predominantly in polymeric form and strongly enriched for λ light chains (∼90% vs ∼51%). This response was accompanied by selective expansion of Gd-IgA1^+^λ^+^ plasma cells, identifying a disease-specific, highly responsive B cell compartment.

**Limitations:** The cytokine combination was selected to reflect key signals present in mucosal immune environments, although in vivo conditions are more complex.

**Conclusions:** Mucosal cytokine signals directly amplify pathogenic B cell responses in IgAN, driving production of polymeric, λ-enriched Gd-IgA1. These findings provide mechanistic validation for therapies targeting the mucosal–systemic axis and offer a biologic explanation for the clinical efficacy of gut-directed corticosteroids and BAFF/APRIL inhibition.

## Introduction

Immunoglobulin A nephropathy (IgAN) is the most common primary glomerulonephritis worldwide and a leading cause of chronic kidney disease progression ^1^. The disease is characterized by mesangial deposition of immune complexes composed of galactose-deficient IgA1 (Gd-IgA1) and anti-glycan autoantibodies, predominantly of the IgG subclass, resulting in formation of nephritogenic immune complexes ^2, 3^. Gd-IgA1 contains *O*-linked glycans deficient in galactose residues within the hinge region, which serve as autoantigenic targets ^4–8^. These complexes, enriched for λ light chains, deposit in the glomerular mesangium, activate complement, and induce mesangial cell proliferation, matrix expansion, and release of proinflammatory and profibrotic mediators, ultimately leading to kidney injury ^9–11^.

Clinically, IgAN is characterized by episodes of macroscopic hematuria frequently associated with infections of the upper respiratory or gastrointestinal tract, implicating mucosal immune activation as a key driver of disease exacerbations ^2, 12–14^. These events are accompanied by increased production of pro-inflammatory cytokines, including interleukin-6 (IL-6), which is elevated in serum and urine of IgAN patients and contributes to disease pathogenesis ^15, 16^. Experimental studies have demonstrated that IL-6 promotes Gd-IgA1 production, while additional cytokines - including IL-21 from follicular helper T cells and the B cell survival factors BAFF and APRIL - drive IgA class switching, plasma cell differentiation, and antibody secretion ^17–19^. Innate immune activation, particularly via Toll-like receptor 9 ^20^, further amplifies these responses through upregulation of IL-6 and APRIL, enhancing pathogenic IgA production ^19^.

Despite these insights, most mechanistic data have been derived from immortalized cell lines or indirect clinical observations ^21, 22^, leaving the response of primary B cells from IgAN patients insufficiently defined. In prior studies, we demonstrated that circulating Gd-IgA1^+^ B cells in IgAN exhibit enrichment for λ light chains and express mucosal homing receptors CCR9, CCR10, and CCR7, suggesting trafficking to mucosal sites ^23, 24^. However, direct functional evidence linking mucosal immune signals to differentiation of these cells into Gd-IgA1-producing plasma cells has been lacking.

In the present study, we evaluated the *in vitro* responses of primary IgA^+^ and Gd-IgA1^+^ B cells from patients with IgAN and healthy controls to a cytokine milieu representing mucosal inflammatory activation. We demonstrate that combined stimulation with IL-6, IL-21, BAFF, and APRIL induces preferential differentiation of IgAN-derived B cells into Gd-IgA1–producing plasma cells, with production of polymeric, λ-enriched Gd-IgA1 closely resembling Gd-IgA1 in nephritogenic immune complexes observed *in vivo*. These findings provide direct mechanistic evidence linking mucosal immune activation to pathogenic Gd-IgA1 production and offer biologic insight into the rationale underlying emerging targeted therapies that modulate the mucosal–systemic axis in IgAN.

## Materials and methods

### Study subjects

Adult patients with IgA nephropathy (IgAN; n=13) were recruited from the Departments of Nephrology, Rheumatology, and Endocrinology at University Hospital Olomouc and the Department of Internal Medicine at University Hospital Ostrava, Czech Republic, between 2020 and 2025. IgAN diagnosis was confirmed by kidney biopsy demonstrating predominant mesangial IgA deposition by immunofluorescence. Baseline clinical data, including age, sex, blood pressure, estimated glomerular filtration rate (eGFR), urinary albumin-to-creatinine ratio, 24-hour proteinuria, hematuria, and use of renin–angiotensin system inhibitors, were obtained from medical records (**Table 1, Suppl. Table S1**). No patient had active macroscopic hematuria at the time of blood collection.

**Table 1.** Clinical and biochemical characteristics of study subjects.

| Characteristic | IgAN | Healthy |
| --- | --- | --- |
| Number of subjects | 13 | 10 |
| Males/Females | 7/6 | 3/7 |
| Age (yr) | 48.8 ± 14.3 | 30.3 ± 5.7 |
| Time since diagnosis (yr) | 16 ± 16 | ND |
| Blood pressure, SBP/DBP mean values (mmHg) | 137/83 | 120/80 |
| eGFR (ml/s per 1.73 m<sup>2</sup>) | 0.98 ± 0.46 | ND |
| Serum creatinine (mmol/l) | 145 ± 103 | ND |
| Serum urea (mmol/l) | 10.7 ± 6.3 | ND |
| Proteinuria (g/d) | 0.4 ± 0.8 | ND |
| Urinary albumin-creatinine ratio (mg/mmol) | 48.1 ± 112.3 | ND |
| Hematuria (Ery/ml) | 20.2 ± 38.0 | ND |
| Disease activity stabilized/ Progression<sup>a</sup> | 12/1 | ND |
| Ongoing glucocorticoid therapy, treated/all | 4/13 | 0/10 |
| ACEi/ARB, treated/all | 11/13 | 0/10 |

Glucocorticoid therapy of IgAN is prednisone. Results are presented as counts for categorical data or mean±SD (apart from blood pressure) for continuous data.
ND, not done; SBP, systolic blood pressure; DBP, diastolic blood pressure; Ery, erythrocytes; ACEi, angiotensin-converting enzyme inhibitor; ARB, angiotensin receptor blocker.
<sup>a</sup>Progression of disease was based on a decrease in eGFR of >10%/year.

Patients with IgA vasculitis were excluded based on systemic manifestations (palpable purpura, arthralgia, abdominal pain). Secondary causes of IgAN, including chronic liver disease, chronic infections, inflammatory conditions, and malignancies, were also excluded.

The control group (n=10) consisted of healthy adult volunteers without evidence of active infection, recruited from the Department of Transfusion Medicine, University Hospital Olomouc. All participants provided written informed consent. The study was approved by the Ethics Committee of the University Hospital and Faculty of Medicine and Dentistry, Palacký University Olomouc (approval number 176/2020).

### Patient and Public Engagement

Patients and the public were not involved in the design, conduct, reporting, or dissemination of this study.

### PBMCs isolation

PBMCs were isolated as previously described ^25^, with several modifications. First, 25 mL of heparinized peripheral blood was diluted with sterile phosphate-buffered saline (PBS, Biosera, Nuaille, France) at a 3:2 ratio and layered onto Histopaque 1077 (Merck Group, Darmstadt, Germany). After centrifugation (400 x g, RT for 30 min), the PBMC layer was collected and washed twice in PBS 200 x g, RT, 20min). PBMCs were successfully isolated from all participants and all samples were analyzed.

### Cytokine Stimulation of PBMCs

Freshly isolated PBMCs were cultured in 24-well culture plates at a density of 1.5 × 10⁶ cells/well in 1 mL of RPMI-1640 medium (Merck Group) supplemented with 10% fetal bovine serum (FBS; Gibco, Waltham, MA, USA) and 1× penicillin-streptomycin (Serana Europe GmgH, Pessin, Germany). Cells were left unstimulated (NS) or exposed to cytokines including IL-2 (10 U/µl), IL-10 (20 µg/mL), IL-17A (160 ng/mL), IL-21 (50 ng/mL), CD40-L (1000 ng/ml), APRIL (100 ng/mL), and BAFF (200 ng/mL) PeproTech [Thermo Fisher Scientific], Cranbury, NJ, USA), and IL-6 (10 ng/mL) R&D Systems, Minneapolis, MN, USA). Following 7-day incubation (37°C, 5% CO_2_), supernatants were collected for immunoglobulin analysis and cells were harvested for flow cytometry.

### Flow Cytometry Staining

Unstimulated or stimulated PBMCs were stained in three steps. The first step consisted in blocking Fc receptors with 10% heat-inactivated human serum in PBS for 10 min at RT, followed by washing the cells and incubation with 41 μg/mL of Texas red-labeled anti-Gd-IgA1 35A12 mAb ^26^ in 5% FBS in PBS for 1 h at 37°C ^24^. In the next step, cells were washed and incubated with fluorophore-labeled mAb specific for cell-surface markers, diluted in 5% FBS in PBS, for 30 min at RT in the dark. All washing steps were performed using 1 ml of PBS followed by centrifugation at 250 x g for 5 min.

Cell-surface molecules were labeled using the following mAbs: anti-CD19 - PE-, FVD – eFluor506 (Thermo Fisher Scientific, Waltham, MA, USA), anti-CD38 - BV650, anti-CD138 - BV711, anti-IgM - BV421, anti-IgD - APC-H7, and anti-CD27 - PE-Cy7 (SONY Biotechnology, San Jose, CA), anti-IgA – FITC (Jackson ImmunoResearch, West Grove, PA, USA) and anti-λ L chain - Pacific Blue (Biolegend, San Diego, CA, USA).

### Flow cytometry analysis

All analyses were performed by the SONY Spectral Analyzer SP6800 (SONY Biotechnology) equipped with blue (488 nm), red (637 nm) and violet (405 nm) lasers. More detailed post-acquisition analyses were done by FlowJo software (BD Biosciences, Franklin Lakes, NJ, USA).

### Determination of Gd-IgA1 and IgA resp. IgG in the serum and tissue culture supernatants

Serum total IgA levels were measured by ELISA as previously described ^27, 28^. Relative Gd-IgA1 concentrations were determined using the KM55 monoclonal antibody (IBL-America), which specifically recognizes terminal N-acetylgalactosamine (GalNAc) residues ^26^. ELISA plates were coated overnight at 4 °C with goat anti-human IgA F(ab′)_2_ (2 µg/mL; Jackson ImmunoResearch), blocked with 1% BSA/PBS/0.05% Tween 20, and incubated with diluted samples. Bound IgA was desialylated ^24^, and Gd-IgA1 was detected using KM55 (0.4 µg/mL), followed by horseradish peroxidase– conjugated goat anti-rat IgG. After development with o-phenylenediamine-H_2_O_2_, absorbance was measured at 490 nm and expressed relative to a Gd-IgA1 myeloma standard.

For IgG quantification, plates were coated with goat anti-human IgG F(ab′)_2_ (2 µg/mL) under similar conditions. After incubation with diluted samples, bound IgG was detected using horseradish peroxidase–conjugated goat anti-human IgG (γ-chain– specific; Merck) diluted 1:6,000. Signal was developed with o-phenylenediamine-H_2_O_2_ and read at 490 nm. IgG concentrations were calculated using a standard curve.

### Detection of kappa/lambda distribution in Gd-IgA1 from supernatants

CBA Beads (BD Biosciences) were conjugated with mouse anti–Gd-IgA1 35A12 mAb according to the manufacturer’s instructions and incubated with supernatants of stimulated PBMCs from IgAN patients and controls for 1 h at 37°C on a shaker. After washing with PBS, anti-kappa – FITC and anti-lambda – PE (Dako, Agilent, Santa Clara, CA, USA) were added for 30 min, at RT, in the dark. After washing with PBS, beads were analyzed by a SONY spectral analyzer (SONY Biotechnology).

### Detection of monomeric and polymeric IgA and Gd-IgA1 from supernatants

Proteins from supernatants were separated by SDS-PAGE electrophoresis under non-reducing conditions using a 5% gel. Separated proteins were transferred onto a PVDF membrane by western blot and analyzed with anti-IgA–HRP (Jackson ImmunoResearch), or anti-Gd-IgA1 and anti–mouse IgG–HRP (Merck Group), respectively.

### Statistics

Normally distributed data were summarized as mean ± standard deviation (SD) and compared via unpaired t-tests or one-way ANOVA with post-hoc Tukey tests between IgAN patients and healthy controls. All statistical analyses were done by GraphPad Prism v.8 software (GraphPad Software, La Jolla, CA, USA).

Bias was minimized through consecutive recruitment and strict eligibility criteria. Standardized laboratory methods ensured consistent measurement across groups. Relevant clinical variables were recorded to address confounding, and controlled *in vitro* conditions reduced biological variability, supporting reliable comparison of immune responses between patients and controls.

No formal calculation was performed concerning the study size, as the exploratory design aimed to detect biologically relevant differences under defined experimental conditions.

## Results

We assessed whether primary IgA^+^ and Gd-IgA1^+^ B cells from peripheral blood of patients with IgA nephropathy (IgAN) ^24^ contribute to the generation of IgA and Gd-IgA1-λ. In contrast to findings from EBV-immortalized cell lines ^29, 30^, primary cells demonstrated minimal responsiveness to IL-6 alone, with only a modest, non-significant increase in Gd-IgA1 secretion (**Suppl. Figure S1**). These results suggest that isolated cytokine stimulation is insufficient to induce clinically relevant IgA production and that a more complex mucosal-like microenvironment is required to drive differentiation of pathogenic B cell populations in IgAN.

To better mimic mucosal immune conditions, we tested in a small cohort of 3 patients and healthy controls multiple cytokine combinations. IL-17, alone or with IL-6, did not significantly affect IgA or Gd-IgA1 production (**Suppl. Figure S2**). The combination of IL-2, IL-6, and IL-10 induced proliferation of Gd-IgA1^+^ cells in IgAN samples but resulted in limited antibody secretion (**Suppl. Figure S3**). Stimulation with IL-6, IL-21, and CD40L preferentially drove differentiation toward IgG-producing cells, with minimal increase in Gd-IgA1 secretion (**Suppl. Figure S4**). Addition of BAFF to IL-6 and IL-21 enhanced IgA, Gd-IgA1, and IgG production (**Suppl. Figure S5**), although enrichment of Gd-IgA1^+^ cells remained limited. Notably, inclusion of APRIL further augmented secretion of IgA, Gd-IgA1, and IgG (**Suppl. Figure S6B**). The combined stimulation with IL-6, IL-21, BAFF, and APRIL showed the strongest effect, promoting expansion of CD19^+^ and Gd-IgA1^+^ cells (**Suppl. Figure S6A**) and maximal IgA and Gd-IgA1 secretion (**Suppl. Figure S6B**).

In summary, the combination of IL-6, IL-21, BAFF, and APRIL displayed the dominant stimulatory effects on Gd-IgA1^+^ cells population and Gd-IgA1 production from PBMC of IgAN patients **(Suppl. Figure S6A)**. Similarly, IL-6, IL-21, BAFF, and APRIL was the more effective combination to increase IgG production by cells from IgAN patients and controls (**Suppl. Figure S6B)**.

### B cells stimulated by combination of IL-6, Il-21, APRIL and BAFF differentiated into CD38^high^ plasmablasts and CD138^high^ plasma cells

To further assess the clinical relevance of these findings, we expanded the cohort to include 13 patients with IgA nephropathy (IgAN) and 10 healthy controls and stimulated PBMCs with a cytokine milieu representative of the mucosal immune environment (IL-6, IL-21, BAFF, and APRIL). The gating strategy is shown in **Suppl. Figure S7**. Under these conditions, B cells preferentially differentiated into antibody-secreting populations (**Figure 1**). Although total viable B cell numbers remained stable, there was a significant increase in plasmablasts (CD38^high^) and plasma cells (CD38^high^ CD138^high^), accompanied by a reduction in naïve and memory B cell subsets (CD38^low^) in both IgAN patients and controls (**Figure 1A, C)**. t-distributed stochastic neighbor embedding (t-SNE) analysis further illustrated these phenotypic shifts, demonstrating distinct redistribution of B cell subsets following stimulation (**Figure 1B**). These findings indicate that mucosal cytokine signals promote differentiation of circulating B cells toward antibody-secreting phenotypes, which represent a key cellular source of pathogenic Gd-IgA1.

**Figure 1.**
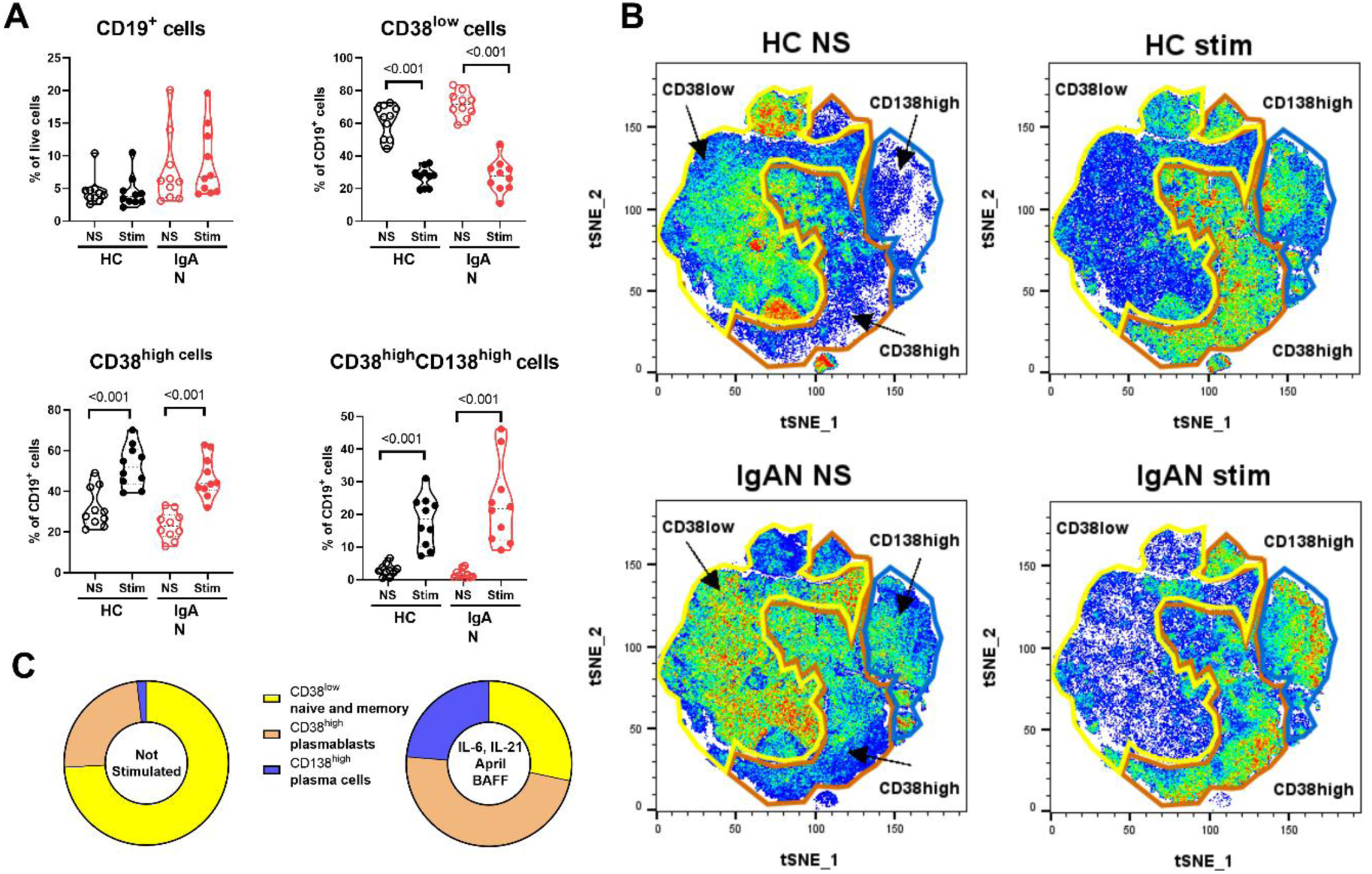
Primary B cells differentiate under mucosal cytokine stimulation into CD38^high^ plasmablasts and CD38^high^ CD138^+^ plasma cells. Cell phenotypes of PBMCs from IgAN patients (*n =* 10) and healthy controls (*n =* 10) before and after stimulation with a combination of IL-6, IL-21, BAFF, and APRIL (Stim) analyzed by flow cytometry **A)**, and characterized by t-SNE (t-distributed Stochastic Neighbor Embedding) visualization **B)**. Differences in cell phenotypes in PBMC of IgAN patients driven by IL-6, IL-21, BAFF, and APRIL are shown **C)**. *P* values were calculated using one-way ANOVA followed by Tukey’s *post-hoc* test. Abbreviations: Gd-IgA1, galactose-deficient IgA1; HC, healthy controls; IgAN, IgA nephropathy; NS, not stimulated; PBMCs, peripheral blood mononuclear cells

### IL-6, IL-21, APRIL, and BAFF cytokine combination significantly stimulated the Gd-IgA1 cells, specifically the cells from IgAN patients

In contrast to minimal changes between IgAN and HC cells in IgA^+^, IgG^+^, and IgM^+^ B cell subpopulations following stimulation with a mucosal inflammatory cytokine milieu (IL-6, IL-21, BAFF, and APRIL), PBMCs from IgAN patients exhibited a significant increase in Gd-IgA1^+^ cells (**Figure 2A**). Immunofluorescence analysis confirmed the presence of IgA^+^ (green) and Gd-IgA1^+^ (magenta) cells (**Figure 2B**), with morphological features consistent with plasmablast and plasma cell differentiation following stimulation. t-SNE analysis further confirmed distinct shifts in immunoglobulin isotype distribution after cytokine exposure (**Figure 2C**).

**Figure 2.**
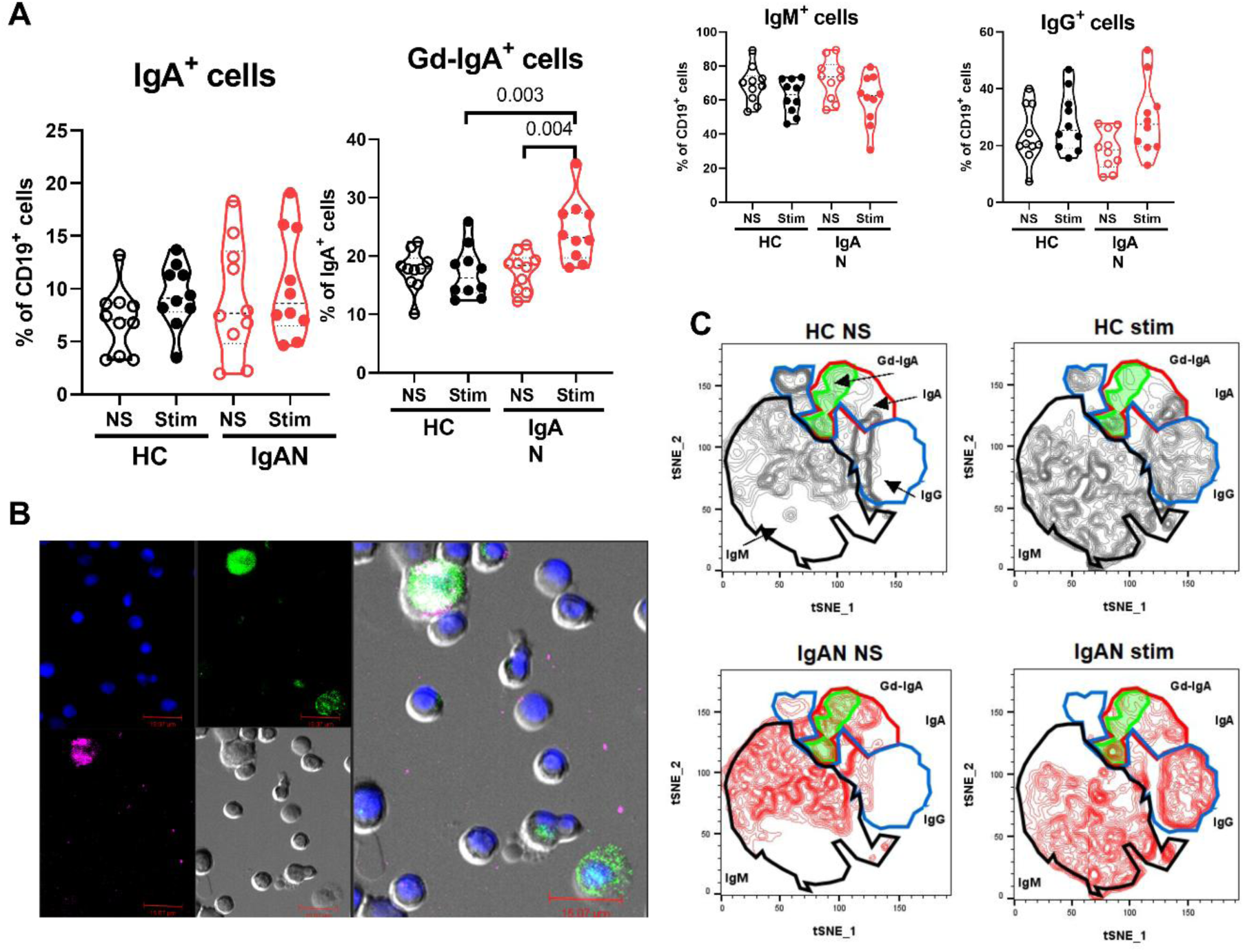
A mucosal environment-mimicking cytokines preferentially promote proliferation of Gd-IgA1^+^ cells in patients with IgAN. **A)** Cell isotypes were detected by flow cytometry before and after stimulation with a combination of IL-6, IL-21, BAFF, and APRIL (Stim) of PBMCs from IgAN patients (*n* = 10) and HC (*n* = 10). **B)** Stimulated cells of IgAN were characterized by fluorescence microscopy (top left: blue (DAPI) - cell nuclei, top right: green (FITC) - IgA, bottom left: magenta (APC) - Gd-IgA1, bottom left: phase contrast, right large panel: overlay). **C)** t-SNE (t-distributed Stochastic Neighbor Embedding) visualization of cell isotype distribution before and after cytokine stimulation of PBMCs from IgAN patients and HC. *P* values were calculated using one-way ANOVA followed by Tukey’s *post-hoc* test. Abbreviations: APC, Allophycocyanin; DAPI, 4′,6-diamidino-2-phenylindole; FITC, fluorescein isothiocyanate; Gd-IgA1, galactose-deficient IgA1; HC, healthy controls; IgAN, IgA nephropathy; NS, not stimulated; PBMCs, peripheral blood mononuclear cells

### Gd-IgA1^+^ cells from IgAN patients had high potential to differentiate into plasma cells

Phenotypic analysis of B cell subsets showed that, although naïve/memory (CD38^low^) and plasmablast (CD38^high^) populations remained largely unchanged, plasma cells (CD38^high^ CD138^high^) were significantly expanded across IgA^+^, IgG^+^, and IgM^+^ compartments in both IgAN patients and controls (**Figure 3A-C**). Notably, enrichment of Gd-IgA1^+^ plasma cells was observed exclusively in IgAN patients (**Figure 3C**). Distribution analysis further confirmed preferential differentiation of Gd-IgA1^+^ cells into the plasma cell compartment (**Figure 3D**).

**Figure 3.**
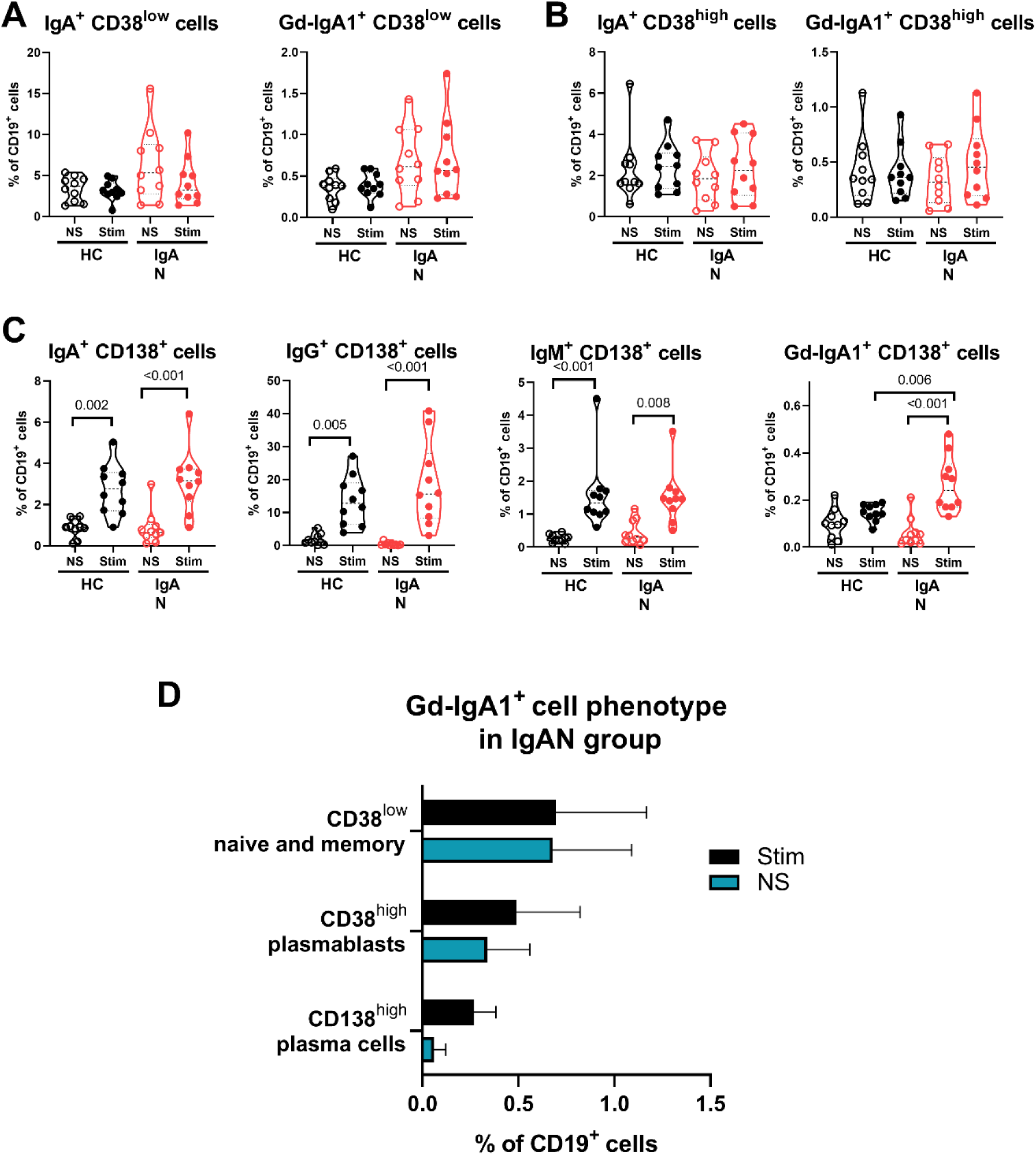
Mucosal cytokine stimulation drives expansion of IgA^+^, IgG^+^, and IgM^+^ plasma cells and selectively increases Gd-IgA1^+^ plasma cells in IgAN patients. **A)** Changes in IgA^+^ and Gd-IgA1^+^ populations of naïve and memory CD38^low^ phenotype in PBMCs of IgAN patients (*n* = 10) and HC (*n* = 10). **B)** Analysis of IgA^+^ and Gd-IgA1^+^ plasma blasts and plasma cells before and after stimulation **C)** Detection of particular isotypes in plasma cells CD138^+^ of stimulated PBMCs of IgAN patients and HCs. **D)** Diagram showing the distribution of Gd-IgA1^+^ cells from the IgAN group based on cell phenotype before and after stimulation. *P* values were calculated using one-way ANOVA followed by Tukey’s *post-hoc* test. Abbreviations: Gd-IgA1, galactose-deficient IgA1; HC, healthy controls; IgAN, IgA nephropathy; NS, not stimulated; PBMCs, peripheral blood mononuclear cells

These findings indicate that B cells from IgAN patients exhibit a selective propensity to differentiate into Gd-IgA1–producing plasma cells under mucosal cytokine stimulation.

### IL-6, IL-21, BAFF, and APRIL stimulated λ chain-positive Gd-IgA1 plasma cells preferentially in IgAN patients

Following stimulation with IL-6, IL-21, BAFF, and APRIL, the proportion of λ light chain-positive cells increased across total CD19^+^, IgA^+^, and IgM^+^ populations (**Figure 4A**), with enhanced λ expression confirmed by fluorescence microscopy (**Figure 4B**). Although enrichment of Gd-IgA1^+^λ^+^ cells was observed in both groups, a significant increase in Gd-IgA1^+^λ^+^ CD138^+^ plasma cells was detected exclusively in IgAN patients (**Figure 4C**). These cells, identified as IgA^+^Gd-IgA1^+^λ^+^CD138^+^ plasma cells (**Figure 4C, D**), likely represent a functionally relevant population with the capacity to produce nephritogenic Gd-IgA1. Their selective expansion in IgAN patients highlights disease-specific sensitivity to mucosal cytokine stimulation and supports their role as a key cellular source of pathogenic Gd-IgA1.

**Figure 4.**
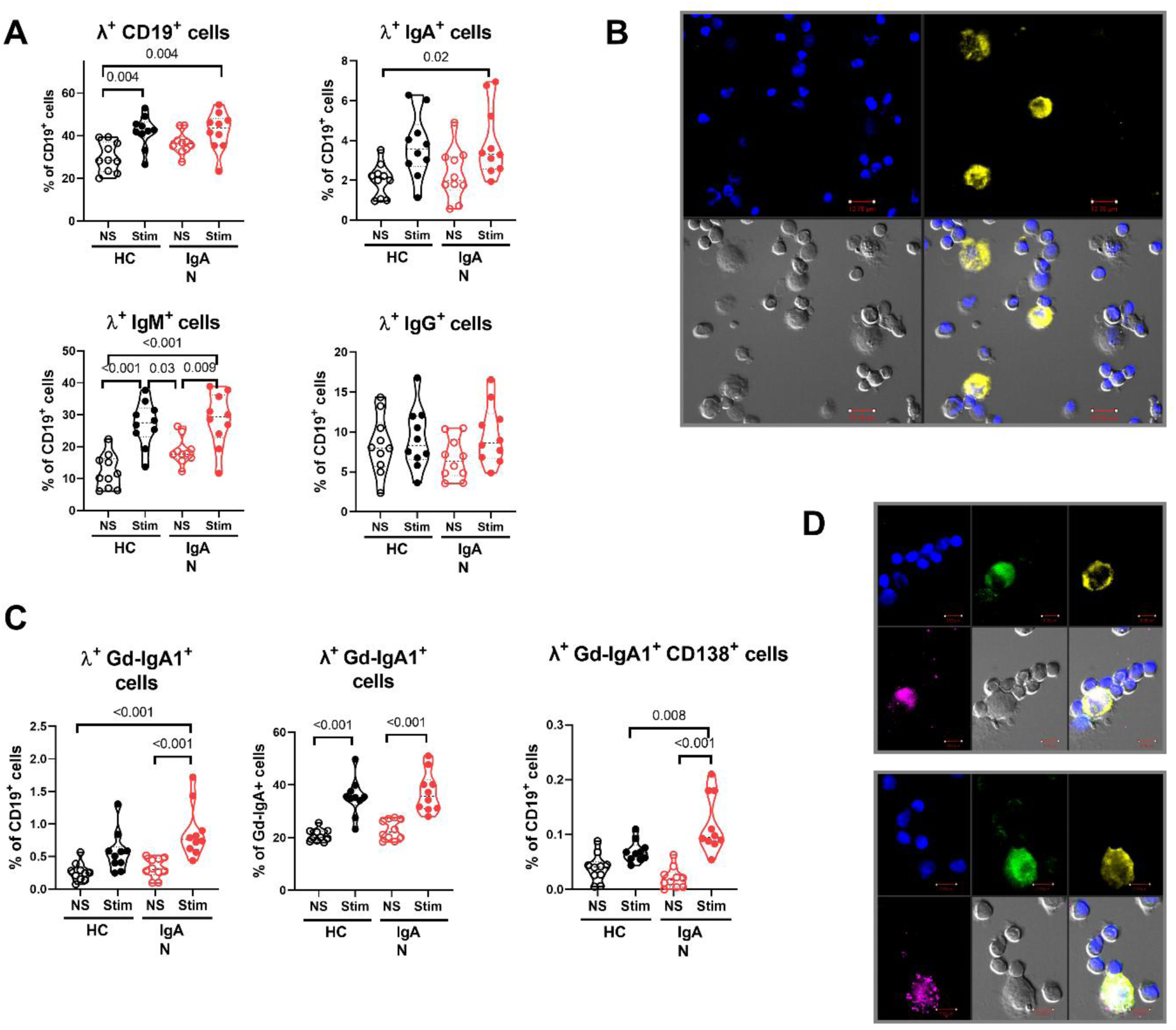
Cytokine stimulation induces selective expansion of Gd-IgA1^+^λ^+^ cells, including Gd-IgA1^+^λ^+^ CD138^+^ plasma cells, in IgA nephropathy. **A)** Lambda light chain expression on CD19^+^, IgA^+^, IgM^+^ and IgG^+^ cells determined by flow cytometry. **B)** Lambda light chain expression on stimulated PBMCs of IgAN patients characterized by fluorescence microscopy. **C)** Surface expression of lambda light chain on Gd-IgA1^+^ cells and on Gd-IgA1^+^ plasma cells determined by flow cytometry. **D)** Colocalization of IgA, Gd-IgA1 and λ light chain on stimulated cells of IgAN patients after IL-6, IL-21, BAFF, and APRIL stimulation by fluorescence microscopy. For all microscopic panels: blue (DAPI) - cell nuclei, green (FITC) - IgA, magenta (APC) - Gd-IgA1, yellow (TRITC) - lambda light chain. *P* values were calculated using one-way ANOVA followed by Tukey’s *post-hoc* test. Abbreviations: APC, Allophycocyanin; DAPI, 4′,6-diamidino-2-phenylindole; FITC, fluorescein isothiocyanate; Gd-IgA1, galactose-deficient IgA1; HC, healthy controls; IgAN, IgA nephropathy; NS, not stimulated; PBMCs, peripheral blood mononuclear cells; Stim, stimulated; TRITC, tetramethylrhodamine isothiocyanate

### Cytokine milieu comprising IL-6, IL-21, BAFF, and APRIL significantly enhances production of IgA, IgG, and polymeric λ-enriched Gd-IgA1 in supernatants of IgAN cells

PBMCs from IgAN patients demonstrated a markedly increased capacity to secrete Gd-IgA1 compared with controls (**Figure 5A**), indicating heightened sensitivity to mucosal cytokine signals. Although a modest increase was also observed in control cells, the response was substantially less pronounced. Importantly, stimulated PBMCs from IgAN patients produced a significantly higher proportion of Gd-IgA1 with λ light chains (approximately 90%) compared with controls (approximately 51%), as determined by CBA bead analysis (**Figure 5B**). Notably, Western blot analysis under non-reducing conditions confirmed that the produced IgA and Gd-IgA1 were predominantly present in polymeric form (**Figure 5C**). This combination of λ light chain enrichment and polymeric structure closely mirrors the composition of pathogenic IgA-containing immune complexes observed in the circulation and mesangial deposits of IgAN patients.

**Figure 5.**
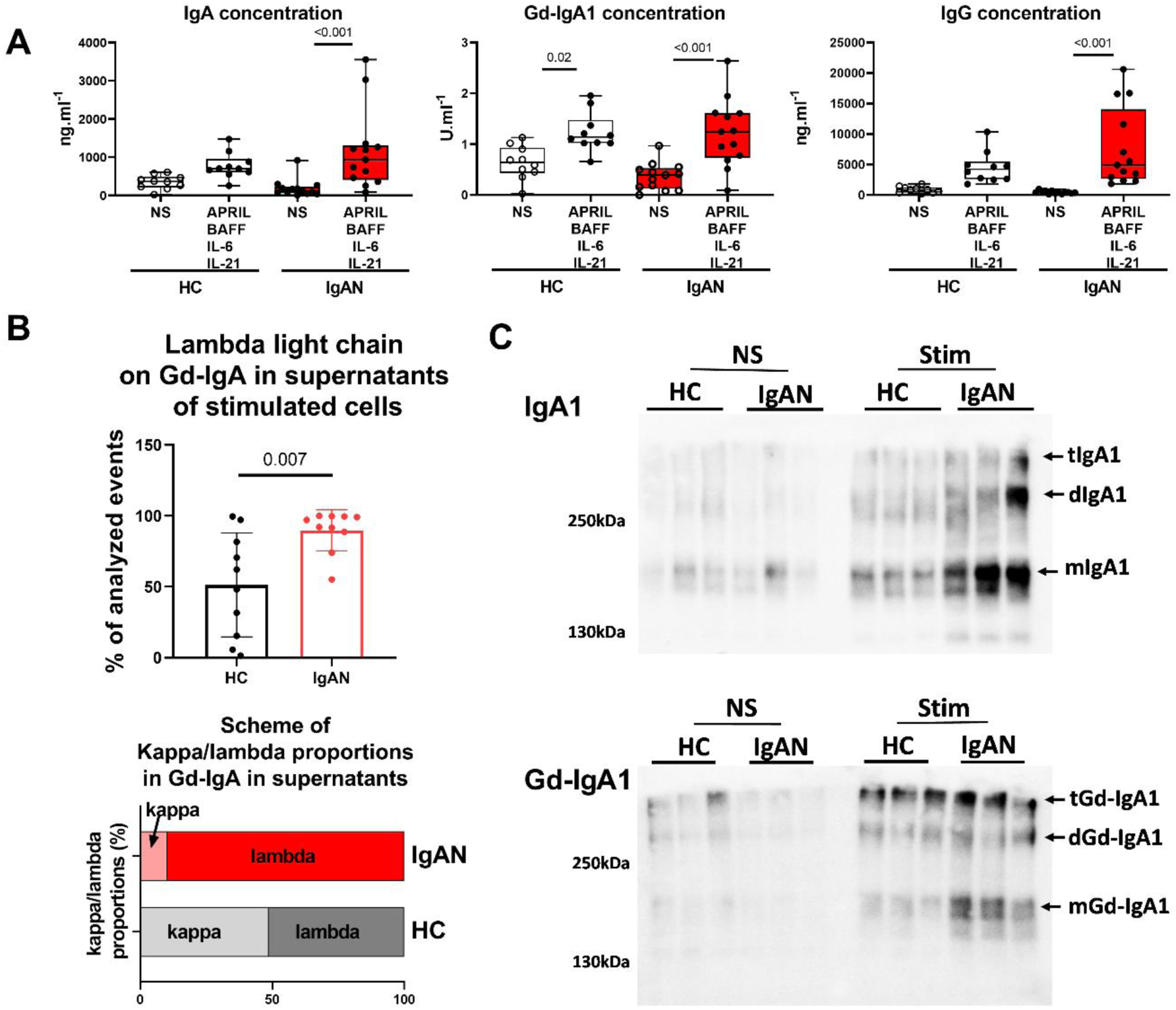
Mucosal cytokine stimulation preferentially enhances production of IgA, λ-enriched polymeric Gd-IgA1 and IgG by PBMCs from IgAN patients. **A)** Levels of IgA, IgG and Gd-IgA1 in supernatants of IL-6 + IL-21 + BAFF + APRIL stimulated cells from HC (*n* = 10) and IgAN patients (*n* = 13) detected by ELISA. **B)** The distribution of λ and κ light chains within Gd-IgA1 molecules in supernatants of stimulated cells from IgAN patients and healthy controls including a schematic representation of κ/λ proportions in both groups. **C**) Detection of monomeric (160kDa) and polymeric (360kDa – dimer) (600kDa – tetramer) structures of IgA and Gd-IgA1 from supernatants from stimulated cells by SDS-PAGE electrophoresis with 5% polyacrylamide gel and western blot using anti-IgA – HRP antibody, and anti-Gd-IgA1 and anti–mouse IgG–HRP staining. P values were calculated using one-way ANOVA followed by Tukey’s post-hoc test. Abbreviations: Gd-IgA1, galactose-deficient IgA1; HC, healthy controls; IgAN, IgA nephropathy; NS, not stimulated; PBMCs, peripheral blood mononuclear cells; Stim, stimulated

## Discussion

Current treatment strategies for IgA nephropathy (IgAN) combine optimized supportive care with emerging targeted therapies, reflecting advances in our understanding of its pathogenesis. Renin–angiotensin system blockade and strict blood pressure control remain foundational, with SGLT2 inhibitors providing additional renoprotective benefit independent of glycemic status ^31, 32^. Systemic corticosteroids can reduce proteinuria in selected patients but are limited by significant toxicity ^33^. More recently, targeted approaches have gained prominence, including gut-directed corticosteroids with targeted release in the distal ileum, which modulate mucosal immune activation within gut-associated lymphoid tissue, and agents inhibiting BAFF and APRIL pathways that influence B cell survival and differentiation ^34, 35^. Complement inhibitors further address downstream inflammatory injury. Collectively, these strategies reflect a shift from nonspecific immunosuppression toward mechanism-based interventions targeting the mucosal–systemic axis and dysregulated B cell responses driving pathogenic Gd-IgA1 production ^32, 33, 36, 37^.

Despite these advances, the cellular mechanisms linking mucosal immune activation to galactose-deficient IgA1 (Gd-IgA1) production remain incompletely defined. The primary site of pathogenic Gd-IgA1 generation has not been conclusively established, although mucosal tissues - particularly the gastrointestinal tract - are strongly implicated based on the origin of IgA responses and the association between mucosal infections and disease exacerbations ^36, 38, 39^. However, direct evidence demonstrating how mucosal signals influence primary circulating B cells to generate nephritogenic IgA has been limited.

In this study, we provide functional evidence supporting the mucosal origin of pathogenic Gd-IgA1. By recreating a mucosal cytokine milieu *in vitro -* comprising IL-6, IL-21, BAFF, and APRIL - we demonstrate that peripheral blood primary B cells from IgAN patients preferentially differentiate into plasma cells producing Gd-IgA1. Importantly, the generated Gd-IgA1 is predominantly polymeric and exhibits a marked λ light chain bias (∼90% in IgAN versus ∼51% in healthy controls), closely resembling the composition of nephritogenic immune complexes detected in circulation and mesangial deposits ^40, 41^. These findings support a mechanistic link between mucosal immune signals and the generation of pathogenic IgA.

Our results extend previous observations on circulating B cell phenotypes in IgAN. We have shown that Gd-IgA1^+^ B cells express mucosal homing receptors, including CCR9 and CCR10, and display λ light chain predominance ^24^. Here, we demonstrate that these cells are not only phenotypically programmed for trafficking to mucosal sites but are also functionally responsive to mucosal cytokine signals. Under these conditions, they undergo preferential differentiation into antibody-secreting cells, supporting a model in which circulating Gd-IgA1^+^ B cells home to mucosal sites, where local inflammatory signals drive their maturation into plasma cells producing nephritogenic Gd-IgA1.

The cytokine combination used in our model reflects key elements of the mucosal immune microenvironment. IL-6, BAFF, APRIL, and IL-21 are produced by epithelial cells, dendritic cells, macrophages, and T cells ^42^. BAFF and APRIL promote IgA class switching and support survival of plasmablasts and plasma cells through TACI and BCMA signaling ^18, 43^, while IL-21 drives terminal B cell differentiation ^44^. IL-6 further supports plasmablast survival and T follicular helper cell function ^45^. Elevated IL-6 levels in IgAN patients and its ability to enhance Gd-IgA1 production underscore its pathogenic relevance ^15, 30^.

These mechanistic insights have direct clinical implications. Compared with systemic corticosteroids, which exert broad immunosuppressive effects, emerging targeted therapies are designed to more specifically interfere with pathways driving Gd-IgA1 overproduction ^36, 46^. Gut-directed budesonide reduces mucosal immune activation at its source within gut-associated lymphoid tissue ^33^, while inhibition of the BAFF/APRIL axis directly disrupts key signals regulating B cell activation, survival, and differentiation into IgA-producing plasma cells ^37^. Notably, BAFF and APRIL signaling has been shown to promote the generation of Gd-IgA1 and its autoantibodies, and targeted blockade of this pathway (e.g., with telitacicept or atacicept) can reduce circulating Gd-IgA1 levels and pathogenic immune complexes ^34, 35, 47^. Our findings provide a biologic explanation for the clinical efficacy of these therapies by demonstrating that mucosal cytokine signals directly modulate primary B cell responses and drive differentiation into Gd-IgA1-producing plasma cells.

In summary, our study bridges a critical gap between clinical observations and cellular mechanisms in IgAN. By demonstrating that mucosal cytokine signals drive differentiation of circulating B cells into plasma cells producing polymeric, λ-enriched Gd-IgA1, we provide direct evidence linking mucosal immune activation to nephritogenic IgA production. These findings reinforce the central role of the mucosal-systemic axis in IgAN and support the rationale for targeted therapeutic strategies.

## Author contributions

K.Z., M.R., and J.M. conceived and designed the study and wrote the manuscript. K.Z., A.C., and P.K. performed the experiments. J.Z., N.P., J.O., and D.G. recruited the patients with IgAN and healthy individuals. K.T. and Y.O. provided the Gd-IgA1-specific mAb 35A12 and actively participated in preparation of the manuscript.

## Data Sharing Statement

The authors confirm that the data supporting the findings of this study are available within the article and its supplementary materials.

