## Supplementary data for "Mucosal Inflammation as a Driver of Gd-IgA1 Production in IgA Nephropathy: Mechanistic Support for Targeted Therapies"

**Supplementary Figure S1. Characterization of the effect of IL-6 on PBMCs from IgAN patients and healthy controls.**

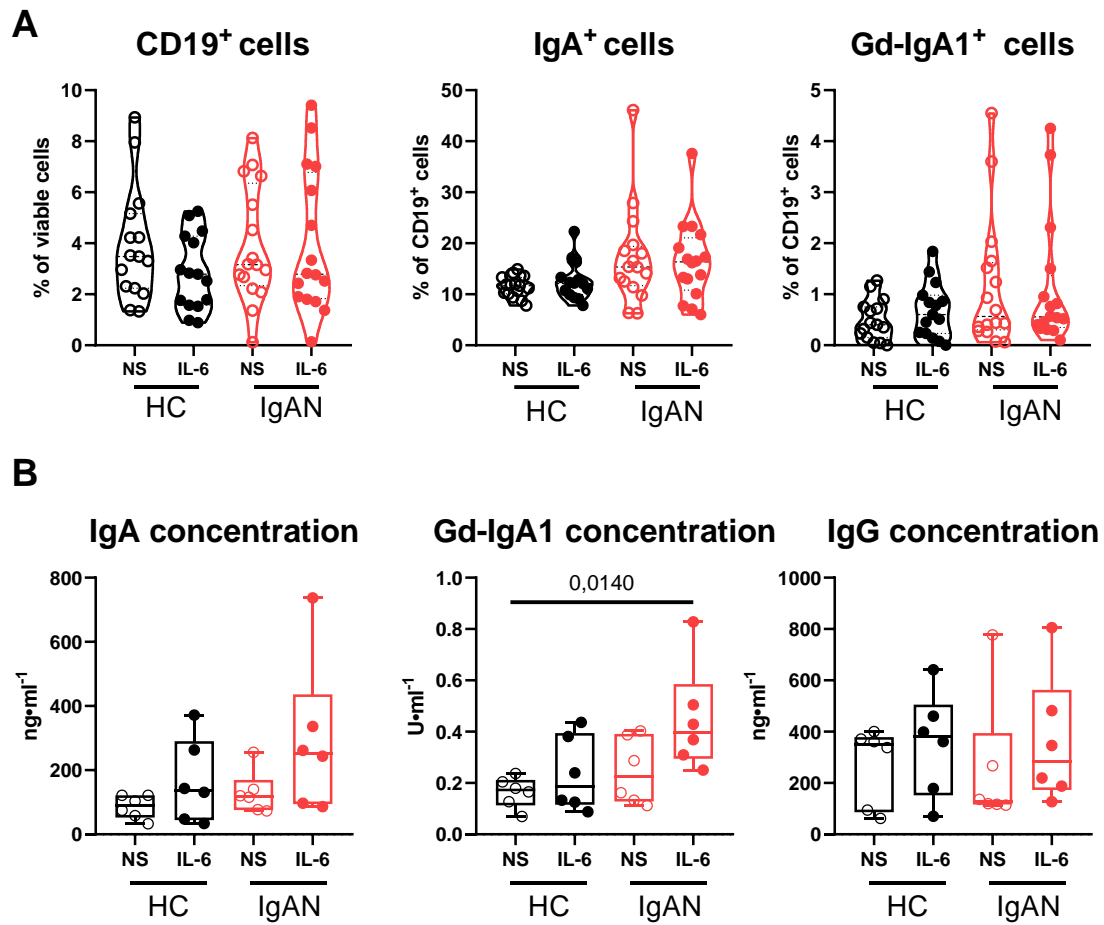

PBMCs from IgAN patients ($n = 15$) and healthy controls ($n = 15$) were stimulated by IL-6 and analyzed by flow cytometry. **A)** No significant changes were detected between unstimulated and IL-6–stimulated cells in the IgAN group or healthy-control cohort in any of the characterized cell populations (CD19<sup>+</sup>, CD19<sup>+</sup> IgA<sup>+</sup> and CD19<sup>+</sup> IgA<sup>+</sup> Gd-IgA1<sup>+</sup> cells). **B)** Additionally, in supernatants from cells of IgAN patients ($n = 6$) and healthy controls ($n = 6$) stimulated by IL-6, no significant increase in IgA, Gd-IgA1, or IgG production was detected. $P$-values were calculated using one-way ANOVA followed by Tukey's *post-hoc* test.

Abbreviations: Gd-IgA1, galactose-deficient IgA1; HC, healthy controls; IgAN, IgA nephropathy; NS, not stimulated; PBMCs, peripheral blood mononuclear cells

**Supplementary Figure S2. Effect of stimulation of PBMCs from IgAN patients with IL-6 and IL-17.**

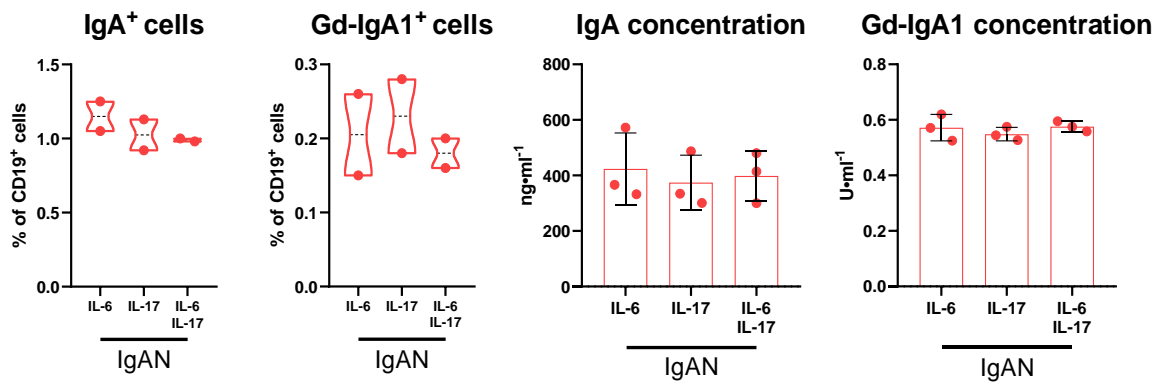

PBMCs from IgAN patients ($n = 2$ or $n = 3$) were stimulated by IL-6, IL-17, or their combination (IL-6 + IL-17) and analyzed by flow cytometry and ELISA. No significant changes were determined in the frequencies of IgA<sup>+</sup> and Gd-IgA1<sup>+</sup> B cells or in the concentrations of IgA and Gd-IgA1 in culture supernatants across stimulation conditions. $P$ values were calculated using one-way ANOVA followed by Tukey's *post-hoc* test.

Abbreviations: Gd-IgA1, galactose-deficient IgA1; IgAN, IgA nephropathy; PBMCs, peripheral blood mononuclear cells

**Supplementary Figure S3. Effect stimulation with IL-2, IL-6 and IL-10 on CD40L-
prestimulated PBMCs from IgAN patients and healthy controls.**

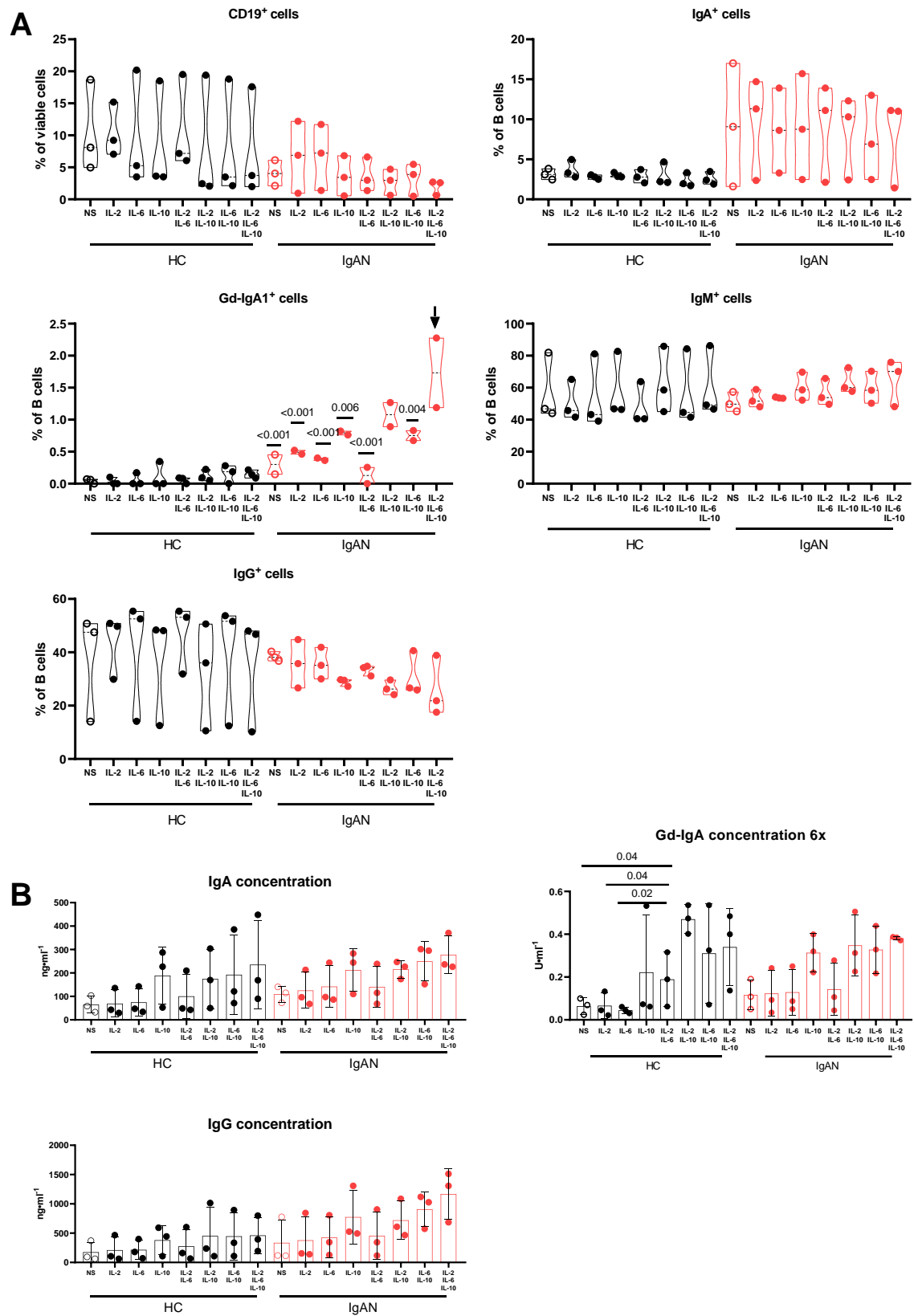

PBMCs from IgAN patients and healthy controls were primed with CD40L and subsequently cultured with a combination of IL-2, IL-6 and IL-10. **A)** Flow cytometry analysis showed that this cytokine combination led to an increase in CD19<sup>+</sup> IgA<sup>+</sup> Gd-IgA1<sup>+</sup> cells, with the increase being significant in the IgAN group. **B)** The increase in Gd-IgA1<sup>+</sup> cell populations did not correspond to higher Gd-IgA1 concentrations in culture supernatants. *P* values were calculated using one-way ANOVA followed by Tukey's *post-hoc* test.

Abbreviations: Gd-IgA1, galactose-deficient IgA1; HC, healthy controls; IgAN, IgA nephropathy; NS, not stimulated; PBMCs, peripheral blood mononuclear cells

#### Supplementary Figure S4. Combination of IL-6, IL-21 and CD40L induces naïve B cells to differentiate preferentially to IgG isotype.

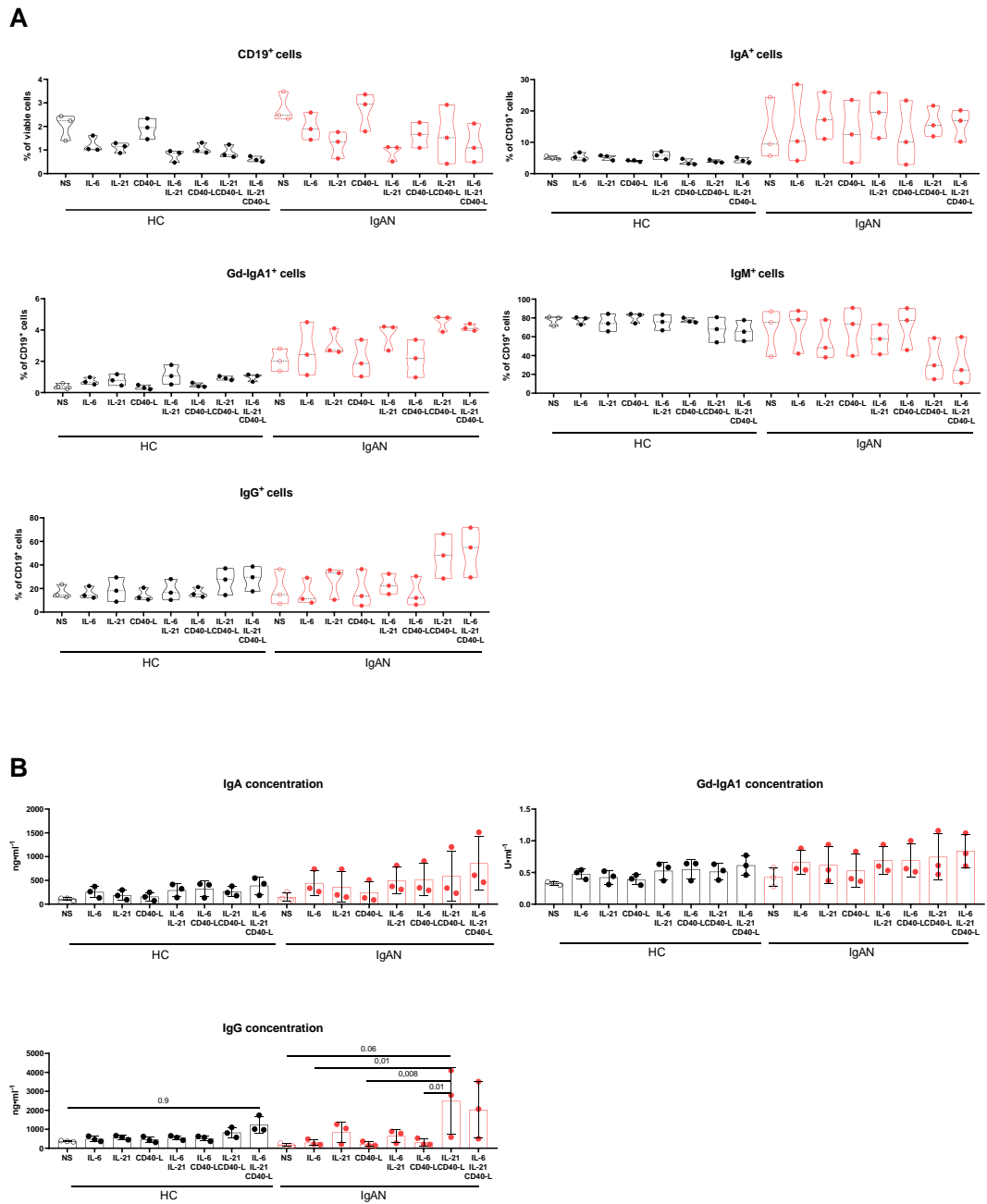

PBMCs from IgAN patients ($n = 3$) and healthy controls ($n = 3$) were stimulated by IL-6, IL-21, CD40L, or their combinations and analyzed by **A**) flow cytometry and **B**) ELISA. **A**) Stimulation by IL-21 + CD40L and IL-6 + IL-21 + CD40L led to a marked increase in CD19<sup>+</sup> IgG<sup>+</sup> cells and a decrease in CD19<sup>+</sup> IgM<sup>+</sup> cells. An increase in CD19<sup>+</sup> IgA<sup>+</sup> Gd-IgA1<sup>+</sup> cells was also observed, particularly in the IgAN group. **B**) IgG

concentrations in culture supernatants significantly increased under these stimulation conditions in IgAN patients. *P* values were calculated using one-way ANOVA followed by Tukey's *post-hoc* test.

Abbreviations: Gd-IgA1, galactose-deficient IgA1; HC, healthy controls; IgAN, IgA nephropathy; NS, not stimulated; PBMCs, peripheral blood mononuclear cells

**Supplementary Figure S5. Effect of IL-6, IL-21 and BAFF on the primary cells from IgAN patients.**

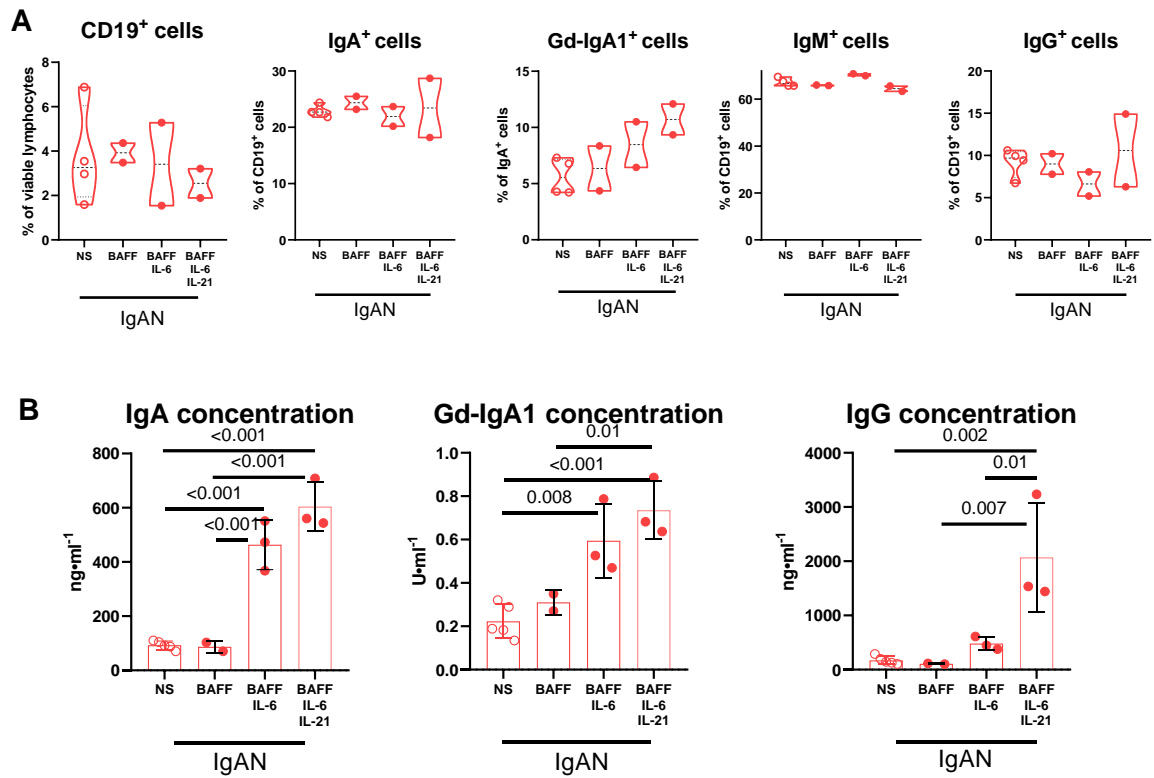

PBMCs from IgAN patients ($n = 2$ or $n = 3$) were stimulated by IL-6, IL-21 and BAFF in various combinations and analyzed by **A**) flow cytometry and **B**) ELISA. **A**) Stimulation by IL-6 + IL-21 + BAFF did not show significant differences in CD19<sup>+</sup>, IgA<sup>+</sup> and Gd-IgA1<sup>+</sup> cells. **B**) In culture supernatants, concentrations of IgA and Gd-IgA1 were significantly increased after IL-6 + BAFF and IL-6 + IL21 + BAFF stimulation. IgG concentration was significantly elevated after IL-6 + IL21+ BAFF stimulation. P values were calculated using one-way ANOVA followed by Tukey's post-hoc test.

Abbreviations: Gd-IgA1, galactose-deficient IgA1; IgAN, IgA nephropathy; NS, not stimulated; PBMCs, peripheral blood mononuclear cells

### Supplementary Figure S6. Summary of different cytokine combination effects on primary cells of IgAN patients and healthy controls.

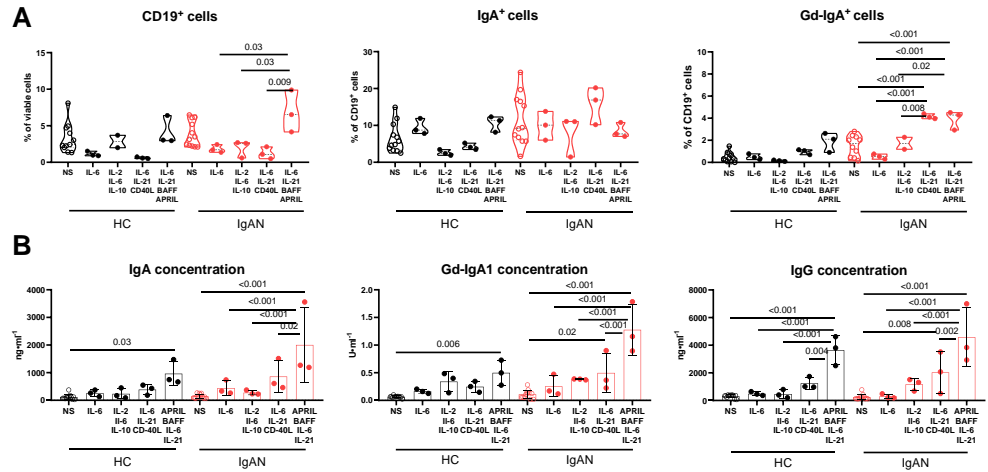

Summary of different cytokine combinations (NS vs. IL-6 vs. IL-2 + IL-6 + IL-10 vs. IL-6 + IL-21 + CD40L vs. IL-6 + IL-21 + BAFF + APRIL) and their influence on A) CD19<sup>+</sup>, IgA<sup>+</sup> and Gd-IgA1<sup>+</sup> cell populations from PBMC of HC ($n = 3$) and IgAN patients ($n = 3$). B) Determination of immunoglobulin production into supernatants of stimulated cells from HC and IgAN patients in different cytokine environments detected by ELISA. P values were calculated using one-way ANOVA followed by Tukey's post-hoc test.

**Supplementary Figure S7. Representative gating of stimulated and unstimulated cells.**

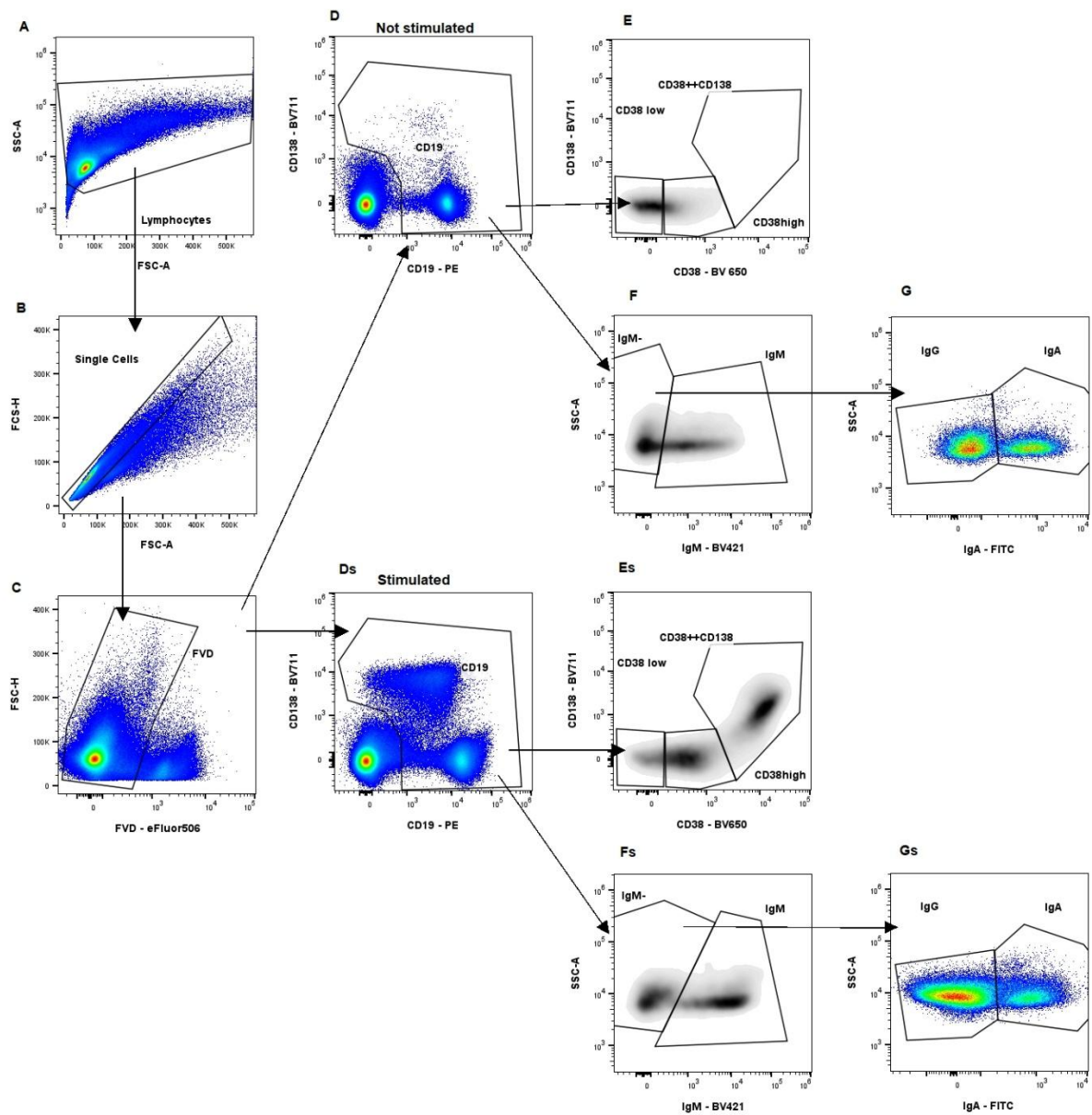

The gating strategy of PBMCs was based (A) on size selection (FSC vs SSC), (B) doublets exclusion (FSC-A vs FSC-H) and (C) dead cells elimination. Next CD19<sup>+</sup> cells were selected (D, Ds) and their phenotypes (E, Es) or isotypes (F, Fs, G, Gs) determination.

**Supplementary Table S1. Individual clinical and biochemical characteristics of study subjects**

|  | Males/Females | Age (yr) | Time since diagnosis (yr) | Blood pressure, SBP/DBP mean values (mmHg) | eGFR (ml/s per 1.73 m<sup>2</sup>) | Serum creatinine (μmol/l) | Serum urea (mmol/l) | Proteinuria (g/d) | Urinary albumin-creatinine ratio (mg/mmol) | Hematuria (Ery/μl) | Disease activity stabilized/progression | Ongoing glucocorticoid therapy (mg/day) | ACEi/ARB, treated/all | MEST |
| --- | --- | --- | --- | --- | --- | --- | --- | --- | --- | --- | --- | --- | --- | --- |
| IgAN1 | M | 72 | 43 | 145/80 | 0.31 | 279 | 20.3 | 0.17 | 10 | 2 | Stab | No | Yes | M1S0E0T0 |
| IgAN2 | F | 69 | 41 | 160/90 | 0.37 | 195 | 12.6 | 0.35 | 32 | 10 | Stab | No | Yes | M1S0E0T0 |
| IgAN3 | M | 29 | 11 | 135/85 | 1.50 | 56 | 6.8 | 0.24 | 10 | 10 | Stab | No | Yes | M1S1E0T0 |
| IgAN4 | M | 50 | 8 | 130/75 | 1.16 | 107 | 10.3 | 0.18 | 1 | 3 | Stab | No | No | M1S0E0T0 |
| IgAN5 | M | 37 | 2 | 125/80 | 0.57 | 206 | 14.2 | 0.41 | 38 | 99 | Stab | 5 | Yes | M1S1E0T1 |
| IgAN6 | F | 68 | 42 | 130/80 | 1.33 | 68 | 6.8 | 0.03 | 1 | 2 | Stab | No | No | M1S0E0T0 |
| IgAN7 | M | 56 | 6 | 150/90 | 0.22 | 408 | 25.9 | 3.07 | 417 | 16 | Stab | 15 | Yes | M1S1E0T1 |
| IgAN8 | M | 43 | 3 | 115/70 | 1.41 | 95 | 5.6 | 0.06 | 3 | 1 | Stab | No | Yes | M1S1E0T0 |
| IgAN9 | F | 54 | 6 | 140/90 | 1.20 | 80 | 9.8 | 0.11 | 19 | 1 | Stab | 10 | Yes | M1S1E0T0 |
| IgAN10 | F | 47 | 16 | 150/90 | 1.50 | 64 | 5.4 | 0.31 | 28 | 2 | Stab | No | Yes | M1S1E0T0 |
| IgAN11 | F | 37 | 4 | 120/80 | 1.00 | 103 | 8.2 | 0.06 | 3 | 3 | Stab | No | Yes | M1S1E0T1 |
| IgAN12 | M | 32 | 1 | 145/85 | 0.93 | 142 | 10.2 | 0.62 | 61 | 111 | Stab | 10 | Yes | M1S1E1T0 |
| IgAN13 | F | 41 | 19 | 130/90 | 1.23 | 82 | 3.5 | 0.06 | 4 | 2 | Stab | No | Yes | M1S0E0T0 |
| HC1 | M | 30 | ND | 110/70 | ND | ND | ND | ND | ND | ND | ND | No | No | ND |
| HC2 | M | 42 | ND | 115/75 | ND | ND | ND | ND | ND | ND | ND | No | No | ND |
| HC3 | F | 27 | ND | 118/78 | ND | ND | ND | ND | ND | ND | ND | No | No | ND |
| HC4 | F | 35 | ND | 122/82 | ND | ND | ND | ND | ND | ND | ND | No | No | ND |
| HC5 | M | 22 | ND | 125/85 | ND | ND | ND | ND | ND | ND | ND | No | No | ND |
| HC6 | F | 35 | ND | 130/88 | ND | ND | ND | ND | ND | ND | ND | No | No | ND |
| HC7 | F | 25 | ND | 115/74 | ND | ND | ND | ND | ND | ND | ND | No | No | ND |
| HC8 | F | 29 | ND | 135/90 | ND | ND | ND | ND | ND | ND | ND | No | No | ND |
| HC9 | F | 28 | ND | 120/80 | ND | ND | ND | ND | ND | ND | ND | No | No | ND |
| HC10 | F | 30 | ND | 110/78 | ND | ND | ND | ND | ND | ND | ND | No | No | ND |

Glucocorticoid therapy of IgAN is prednisone.

ND, not done; Stab, stabilized.
